# Complex Reticulation in Backbone Subfamily Relationships in Leguminosae

**DOI:** 10.1101/2024.07.12.603319

**Authors:** Jacob Stai, Warren Cardinal-McTeague, Anne Bruneau, Steven Cannon

## Abstract

Contradictory lines of evidence have made it difficult to resolve the phylogenetic history of the legume diversification era; this is true for the backbone topology, and for the number and timing of whole genome duplications (WGDs). By analyzing the transcriptomic data for 473 gene families in 76 species covering all six accepted legume subfamilies, we assessed the phylogenetic relationships of the legume backbone and uncovered evidence of independent whole genome duplications in each of the six legume subfamilies. Three subfamilies — Cercidoideae, Dialioideae, and Caesalpinioideae — bear evidence of an allopolyploid duplication pattern suggestive of ancient hybridization. In Cercidoideae and Dialioideae, the hybridization appears to be within-subfamily, with the genera *Cercis* and *Poeppigia* apparently unduplicated descendants of one of the parental lineages; in Caesalpinioideae, the hybridization appears to involve a member of the Papilionoideae lineage, and some other lineage, potentially extinct. Three independent lines of evidence, consisting of a concatenated superalignment, concordance factor analysis of the set of gene family alignments and topologies, and direct inference of reticulation events via maximum pseudo-likelihood implemented by PhyloNet, converged on a single backbone hypothesis and the above hypotheses of reticulate evolution.

**Significance Statement:** In a hybridization event, genes that have already been evolving separately for potentially millions of years become sister chromosomes, yet remain related to one another not at the moment of hybridization, but at the speciation node of the hybrid’s parents. Methodologies based on counts of bursts of duplicated genes, can therefore be fundamentally vulnerable to incorrect and contradictory conclusions about the number and timing of WGD events, unless interpreted carefully and in combination with data from gene trees discordant with the consensus backbone. Our assessment of the legume backbone in that light, resolves previous contradictory findings by concluding that three legume subfamilies are allopolyploid relative to the ur-legume.

---

The third-largest plant family in terms of species number, and the source of billions of tons of food both for humans and livestock, Leguminosae is perhaps best known for its fertilizer-producing root nodule, which enables the fixation of atmospheric nitrogen via a symbiotic relationship with anaerobic soil bacteria. Efficiently stewarding the many genetic and economic resources of legumes requires accurate understanding of the family’s evolutionary history. Phylogenetic analysis of Leguminosae has established the monophyly of six legume subfamilies: Papilionoideae, Caesalpinioideae, Dialioideae, Cercidoideae, Detarioideae, and a monospecific subfamily for the species *Duparquetia orchidacea* Baill. (1)

The legume backbone relationships have been difficult to resolve, with the most recent publication of the Legume Phylogeny Working Group leaving it partly polytomous (1). A few of the most recent studies with good sampling coverage across legume subfamilies had found agreement on a common legume backbone phylogeny (2–4): ((((Pap., Cae.), Dia.), *Dup*.), (Cer., Det.)), but alternative models that vary slightly from this have continued to appear in the literature: a model with Cercidoideae sister to the remaining five (5, 6); a model with Dialioideae sister to Caesalpinioideae (7); and most recently a model with Duparquetioideae sister to Dialioideae (8). Additionally, although studies at the broadest level agree that the evolutionary history of the legumes has been marked by whole genome duplication (WGD), disagreement continues regarding the precise number and timing of these duplications. Koenen et al. (3) found, on the basis of a gene count methodology, independent autopolyploidy at the base of each of Papilionoideae and Detarioideae, in agreement with previous research (9), but suggested that the Papilionoideae WGD was most likely preceded by another allopolyploid WGD shared with Caesalpinioideae and Dialioideae, discarding previous hypotheses of a Caesalpinioideae-specific WGD. This latter allopolyploid WGD hypothesis was derived from the Gene-tree Reconciliation Algorithm with MUL-trees for Polyploid Analysis (GRAMPA) (10). Zhao et al. (4) were in agreement with Koenen et al. (3) regarding the hypothesis that multiple rounds of WGD affect Papilionoideae; however, on the basis of counts of “bursts” of duplicated genes at various nodes, they placed the other WGD affecting Papilionoideae at the crown node of all legumes. This hypothesis of a WGD affecting all legumes was also in agreement with previous research (11). Zhao et al. (4), however, did not discard hypotheses of a Caesalpinioideae-specific WGD; indeed, they concluded that each legume subfamily (except Duparquetioideae) bore evidence of independent WGDs, subtending the crown node of each, in addition to a pan-legume WGD, with most subfamilies in their model having undergone two successive rounds of WGD.

Complicating all these assessments is the parallel observation made in multiple recent papers (5, 6, 12) that the genus *Cercis* L. is diploid, bearing no evidence of having undergone any WGD during the legume diversification era, much less two successive rounds, despite descending from the common ancestors of Leguminosae and Cercidoideae. Although the remaining Cercidoideae bear evidence of WGD, Stai et al. (5) proposed that this clade (i.e., all Cercidoideae genera sister to *Cercis*) is an allopolyploid descendant of two parents on or sister to the *Cercis* lineage, based substantially on analysis of Ks plots. Sinou et al. (12) reached the same conclusion in their analysis of the transcription factor and regulator of floral symmetry CYCLOIDEA. Zhong et al. (6) reached the same conclusion in their analysis of a chromosomal-level genome assembly of *Bauhinia variegata* L. While Koenen et al. (3) settled on an allopolyploid backbone model that could theoretically accommodate an observation of *Cercis* non-polyploidy, no model that can accommodate such an observation can contain the pan-legume WGD proposed by Zhao et al. (4).

Allopolyploidy arises from the inheritance of multiple complete copies of the genome from distinguishable parental lineages. Genes which were once orthologs to each other in the parental lineages, become homeologs post-allopolyploidization. Two moments mark the genomic history of hybrid species; the moment of the parents’ speciation from one another, and the moment of hybrid origin. Polyploidy (allopolyploidy included) then initiates a process of post-polyploid diploidization (13), with chromosomal rearrangements typically resulting in reductions in chromosome number (descending dysploidy), and with dysploidal variations frequently contributing to cladogenesis by effecting reproductive isolation. Chromosomal rearrangements of this nature can result in homeologous recombination (14, 15) leading to paralog substitution, up to and including the substitution of entire paralogous chromosomes (16). As Mason and Wendel write (17), the consequences of these homeologous exchanges are numerous: changing “allele dosage, genomewide methylation patterns and downstream phenotypes”; inducing both speciation and genome stabilization; and potentially “lead[ing] to the production of genomes which appear to be a mix of autopolyploid and allopolyploid segments, sometimes termed ‘segmental allopolyploids.’” In short, due to the large number of genes with distinct origins, allopolyploidy creates a structured source of persistent gene-tree species-tree discordance. In this analysis, we integrate several lines of evidence — a concatenated superalignment, concordance factor analysis of the set of gene family alignments and topologies, and direct inference of reticulation events via the program PhyloNet — to address persistent issues resolving the topology of the legume backbone.

## Results

### Backbone Species Trees Accord With Previous Studies

The single-copy consensus species tree (Fig. 1) accords with previous studies (2–4), finding Papilionoideae (PAP) and Caesalpinioideae (CAE) as sisters to one another; Dialioideae (DIA) as sister to that pair; *Duparquetia* (DUP) as sister to those three (99% bootstrap support); and Cercidoideae (CER) and Detarioideae (DET) as sister to one another with low (41%) bootstrap support, sister to the other subfamilies.

**Fig. 1.**
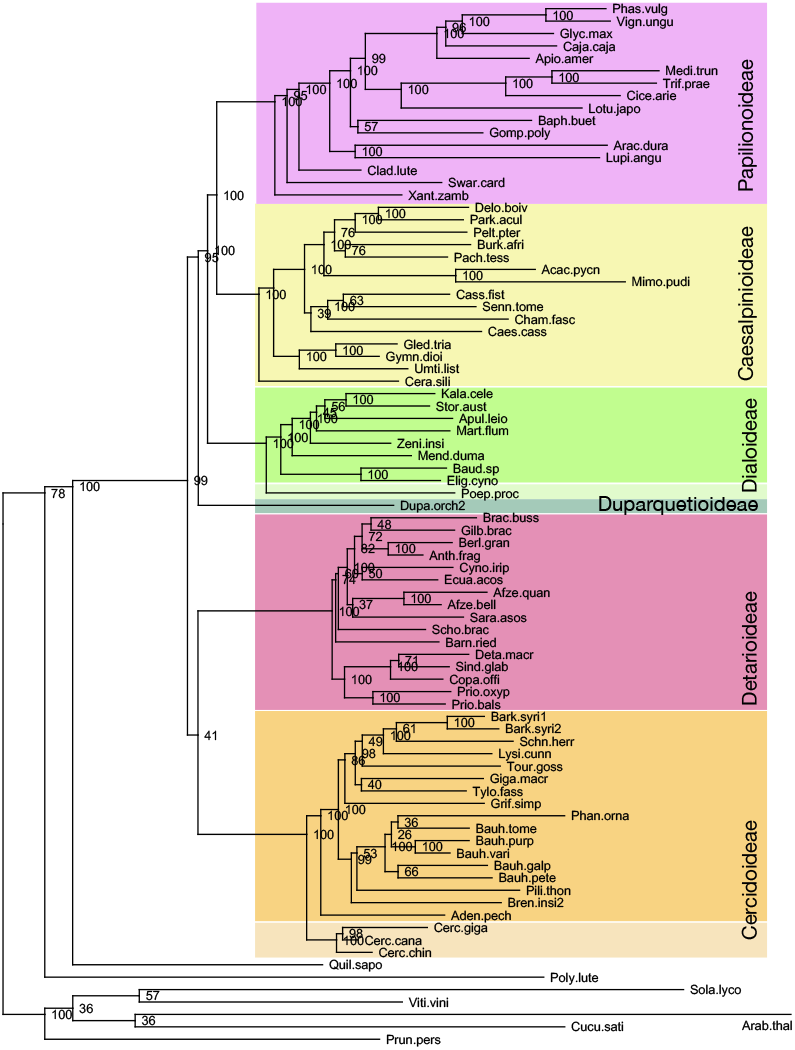
Consensus species tree. This maximum likelihood phylogeny was calculated from 50 gene families with high completeness and sequence representation for the selected species. From each gene family, one sequence was selected (on the basis of sequence length) for inclusion in a supermatrix that was used for phylogeny reconstruction. This reconstruction assumes bifurcation at each node rather than reticulate relationships, in contrast with other analyses presented in this paper. The unduplicated genera *Cercis* and *Poeppigia* are highlighted in lighter tints of orange and green.

However, as in previous studies, complications arose in determining the placement of whole genome duplications. Fig. 2 shows the best-scoring maximum likelihood tree from our duplication clade analysis. In retaining two gene copies, the tree displays both species and hypothetical pseudo-subgenome relationships at once. Bootstrap support is relatively high at the crown nodes of some individual subfamilies — e.g. PAP, 92; CER, 99; DET, 98 — but is often low throughout the rest of the tree, including at the remaining backbone nodes. Caesalpinioideae duplication clades are not sister to one another. One is affiliated with Papilionoideae (boostrap support: 69). The other is affiliated with Dialioideae (though with low bootstrap support, 47; note that a Caesalpinioideae-Dialioideae affiliation was also found in Baker et al. (7)). Dialioideae duplication clades are not sister to one another either, with one clade more closely affiliated to a Caesalpinioideae clade. The two *Duparquetia* sequences appear as sister to the rest of the legumes, but along a backbone with support low enough to be effectively polytomous. Within Dialioideae, the single *Poeppigia* sequence nests within the A subgenome clade (sister to the remaining genera except for *Martiodendron* Gleason and *Zenia* Chun). This would be consistent with an allopolyploid model of Dialioideae evolution with *Poeppigia* a potential candidate for one of the Dialioideae subgenome progenitors. Within Cercidoideae, our combined species-subgenome tree is also consistent with an allopolyploid model, in which the genus *Cercis* is a direct unduplicated descendant of the parent of a subgenome of the remaining Cercidoideae. A and B clades of non-*Cercis* Cercidoideae appeared as sister to one another at a higher frequency than occurred in DIA, such that Fig. 2 offers less support for a Cercidoideae allopolyploid model than a DIA allopolyploid model, in contrast to the evidence from counts of duplicated genes.

**Fig. 2.**
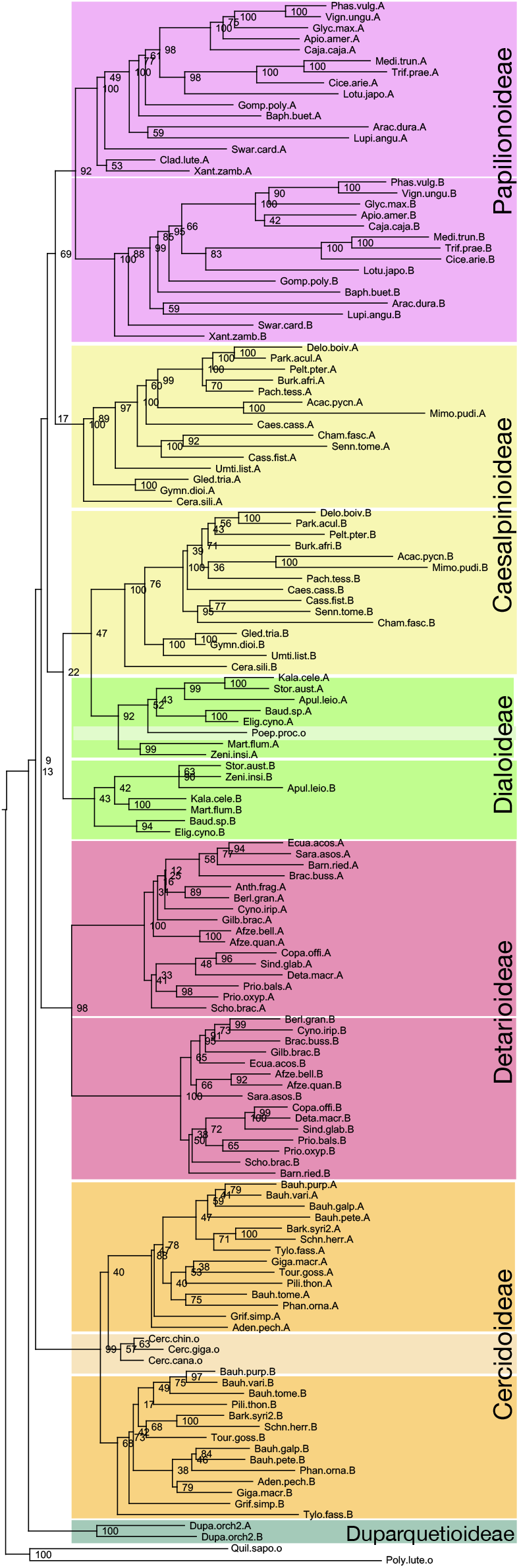
Consensus species-and-whole-genome-duplication tree. This maximum likelihood phylogeny was calculated from 36 gene families with high completeness and sequence representation for the selected species, with each legume sequence being labeled as A or B on the basis of duplication clades observed in previously constructed gene family phylogenies. (See Methods for description of scoring and labeling protocols). From each gene family with labeled genes, up to two genes (A and B) were retained from each species and concatenated into a supermatrix that was used for phylogeny construction.

### *Cercis* and *Poeppigia* Sequence Counts Offer No Evidence of Either Species Having Undergone a WGD Relative to the Legume Ancestor

Counts of recovered genes per species, in each family of our set can be found in Supplemental Table S2. Two species were striking exceptions to baseline rates of retained gene duplicates, in our set of families selected for duplicate retention: *Poeppigia procera* (Poepp. ex Spreng.) C.Presl (DIA) and *Cercis canadensis* L. (CER). Baseline rates ranged from 26.1% for PAP, to 7.8% for CER; but for *Poeppigia*, we found no instances of duplicated genes across the 50 selected gene families (and 3.5% for *Poeppigia* across all gene families; compare to 16.2% in select and 8.8% across all, for the rest of Dialioideae). For *Cercis*, the rate was 1.3% (only increasing to 1.4% when considering all gene families; compared to 7.8% in select and 6.8% across all, for the rest of Cercidoideae). In our visual assessment of those few families (15) in which multiple genes for a single *Cercis* or *Poeppigia* species appeared, we found no clear example of a gene tree individually interpretable as indicating inclusion of either *Cercis* or *Poeppigia* in a WGD event subtending their respective subfamilies.

Both *Cercis* and *Poeppigia* are sister to the other members of their respective legume subfamilies (Fig. 1). Species occupying equivalent phylogenetic positions, were not found to display a similar lack of duplications. *Ceratonia* L. (Caesalpinioideae), was duplicated in 10% of families in which it appeared; *Xanthocercis* Baill. (Papilionoideae), 19.7%. The same was true of the clades sister to *Cercis* and *Poeppigia. Adenolobus* (Harv. ex Benth.) Torre & Hillc., sister to the rest of the non-*Cercis* in Cercidoideae, was present in duplicate in 22% of the gene families; for a clade of *Eligmocarpus* Capuron and *Baudouinia* Baill., sister to the rest of the non-*Poeppigia* Dialioideae, at least one species was present in duplicate 40.7% of the time. Although we only have one of the two *Poeppigia* species in our dataset, for *Cercis*, of the six species sequenced, all were present in duplicate much less often than any other early diverging genera. Sequence counts thus offer no evidence that either *Cercis* or *Poeppigia* have undergone a WGD since the time of the legume ancestor, unlike the other early-diverging genera that we assessed.

In analyzing the topologies of 439 of our 473 gene families (Table 1; families with poor recovery of single-subfamily clades were filtered out), for all gene families with duplication clades for a particular subfamily, the duplication clades resolved as sister to one another with the following frequencies: PAP, 75%; CAE, 16%; DIA, 77%; DUP, 79%; CER, 90%; and DET, 68%. However, for DIA, in which *Poeppigia* is unduplicated, in 68% of the total clades, *Poeppigia* was more closely related to one clade of non-*Poeppigia* DIA, than to the other. Likewise, clades of *Cercis* genes are resolved as more closely related to one clade of non-*Cercis* CER, than to the other, 64% of the time. The same pattern is found in Fig. 2. Regarding DUP, 79% of genes present in a single tree in duplicate resolved as sister to one another; however, the duplication was typically on a long branch close to the legume backbone, and split patterns were common.

**Table 1.** Monophyly rates for duplication clades of genes from the six legume subfamilies.

| Subfamily | PAP | CAE | DIA | DUP | CER | DET |
| --- | --- | --- | --- | --- | --- | --- |
| % Monophyletic | 75 | 16 | 77 | 79 | 89 | 68 |
| (excl. *Cercis* & *Poeppegia*) | — | — | 7 | — | 24 | — |

Clades of non-*Cercis* Cercidoideae and non-*Poeppegia* Dialioideae often were not sister to one another.

It is important to reiterate that for CAE, of all gene families bearing evidence of any CAE whole genome duplication, in the vast majority (84%), the two duplication clades did not resolve as sister to one another. No other legume subfamily was found to be marked as strongly by intersubfamilial genetree species-tree discordance. Out of the 335 instances of CAE clades appearing as sister to a specific other subfamily, modally, 146 times, PAP was that sister; this occurred about twice as often as DIA, the second-most-frequent sister lineage to CAE with 78 occurrences.

### Concordance Factors Agree with the Expected Topology, except regarding *Duparquetia*

In our analysis of gene concordance factors (gCFs; Fig. 3a), we identified two pairs of subfamilies that were most likely to resolve as sister to one another in our gene trees, relative to other pairs: PAPCAE and CER-DET. This is expected, given the consensus backbone relationships. Likewise, DIA and DUP were generally more likely to resolve along the legume backbone than to resolve as sister to a single specific other legume subfamily. Inference of the backbone subfamily topology by neighbor-joining, using gCFs as a distance matrix between subfamilies, reconstructed the expected topology of ((((PAP, CAE), DIA), DUP), (CER, DET)).

**Fig. 3.**
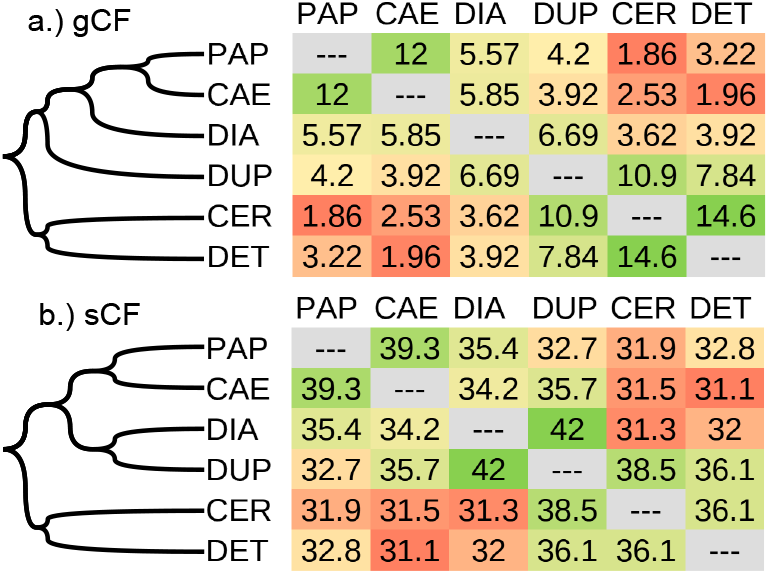
Concordance factor analysis. All 945 possible backbone topologies were scored against 219 low-copy gene family topologies and associated alignments to generate gene and site concordance factors (gCFs / sCFs) via IQTree. The highest gCF / sCF value for each terminal-clade hypothesis was identified. The backbone topology inferred from the resulting gCF / sCF distance matrix via a neighbor-joining algorithm is displayed at left.

However, concordance factor analyses showed variation in the placement of *Duparquetia* within our set of gene trees. Although the most common position for *Duparquetia* genes was subtending a clade of ((PAP, CAE), DIA), when considering only subfamily-pair hypotheses, *Duparquetia* genes occupied more frequently a position subtending CER, than subtending any of the three subfamilies. Furthermore, sCFs (Fig. 3b) indicated that DIA and DUP were the most likely of any two subfamilies to be identical to one another at any particular alignment site. Consequently, the backbone subfamily topology inferred by neighbor-joining using sCF-based distance matrices (Fig. 3b), was (((PAP, CAE), (DIA, DUP)), (CER, DET)). This model (found previously in Zuntini et al. (8)) was inconsistent with the gCF-based topology (Fig. 3a) with respect to the placement of *Duparquetia*, but was otherwise in agreement regarding all the other subfamilies.

### Caesalpinioideae Allopolyploidy Inferred by Maximum Pseudo-Likelihood in Reticulate Network

PhyloNet was constrained to infer by maximum pseudo-likelihood a reticulate phylogenetic network consistent with a topology of ((((PAP, CAE), DIA), DUP), (CER, DET)), with reticulation events affecting the following three or four subfamilies: CAE, DIA, CER, and then optionally DUP (Fig. 4). *Cercis* and *Poeppigia* were treated separately from their respective subfamilies. The non-*Cercis* CER and the non-*Poeppigia* DIA were assessed as originating in a hybridization event, with *Cercis* and *Poeppigia* descendant from one ancestor, and a sister lineage serving as the second parent. For CAE, the likeliest hypothesis of reticulation was found to involve a hybrid origin, with a member of or near-sister of the PAP lineage serving as one ancestral parent, with a lost lineage ancestral to all legumes serving as the other parent (Fig. 4a, b). Results for the other three subfamilies were the same even when DUP as an additional hybrid lineage was included (Fig. 4b). The likeliest hypothesis of reticulation involving *Duparquetia* assigned it a hybrid origin. One parent was a member of or near-sister of the lineage which gave rise to PAP, CAE, and DIA, and the other was a lost lineage ancestral to all legumes (in accordance with the phylogenetic position of DUP in Fig. 2). Notably, the three-reticulation model affecting CAE, the non-*Poeppigia* DIA, and the non-*Cercis* CER, was found by PhyloNet to be not only the likeliest three-reticulation model (Fig. 4a), but also to be likelier than any model involving two reticulations and unspecified for allopolyploid subfamilies (Fig. 4c). Our four-reticulation model including DUP (Fig. 4b), was also found to be likelier than the likeliest three-reticulation hypothesis unspecified for allopolyploid subfamilies (Fig. 4d).

**Fig. 4.**
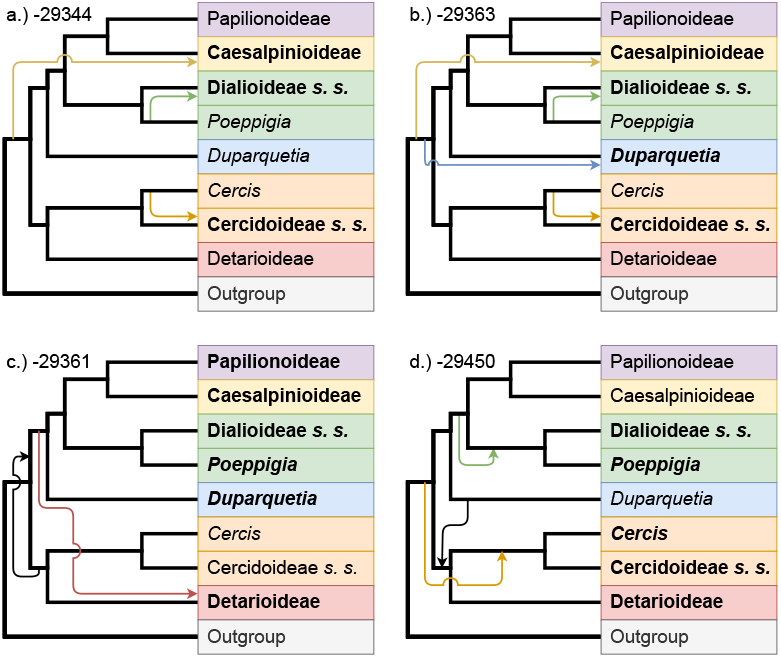
Backbone subfamily networks inferred using maximum pseudo-likelihood. Networks were calculated using PhyloNet (Yu and Nakhleh 2015), from 473 gene families whose backbone topologies were manually scored. A and B duplication clades were mapped explicitly to their common subfamilial taxa. The starting backbone topology was fixed to the consensus backbone. Taxa involved in the reticulation event are indicated in bold. The likeliest additional reticulation event hypotheses were returned for a.) a 3-reticulation event model, with reticulation events affecting the subfamilies Caesalpinioideae, Dialioideae, and Cercidoideae; b.) a 4-reticulation event model of legume backbone evolution, affecting the subfamilies above plus Duparquetioideae; the likeliest c.) 2-reticulation and d.) 3-reticulation models affecting an unspecified set of subfamilies.

### A Working Model of the Legume Backbone with Duplications

Based on the concatenated-supermatrix phylogenies (Figs. 1, 2), the concordance relationships (Fig. 3), and the reticulate networks (Fig. 4) in the analyses described above, and the large body of literature on legume phylogeny, our working model of the legume backbone is shown in Fig. 5.

**Fig. 5.**
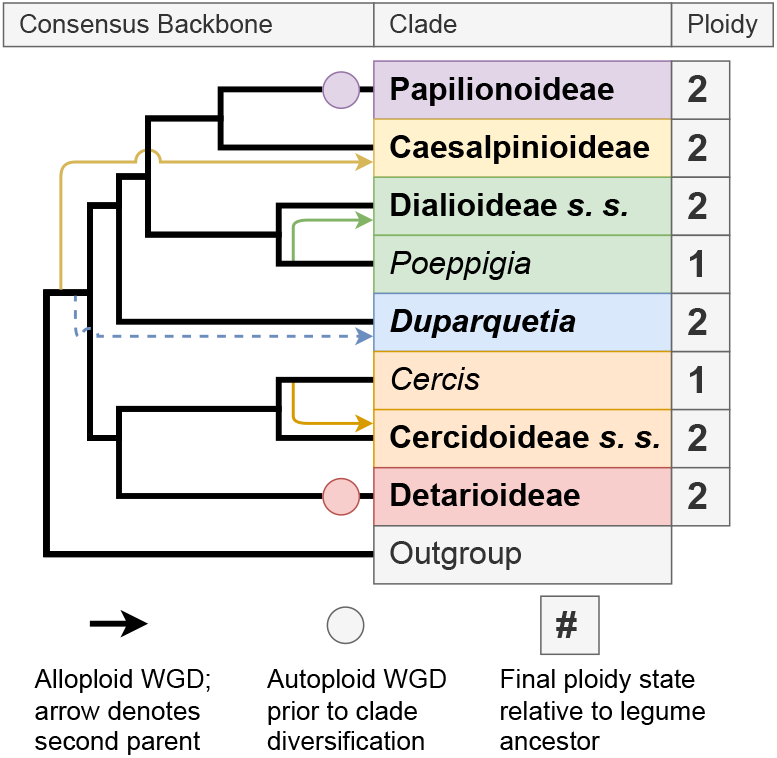
A consensus phylogenetic model of the legume subfamily backbone relationships, based on our results and prior literature. Inferred duplication state from counts of duplicated genes is marked at right; polyploid subfamilies are thus marked in bold. Autoploid WGDs are marked as circles below the relevant subfamilies. Putative allopolyploidies are marked as arrows into the relevant subfamily, with the position of the base of the arrow positioned to reflect the likeliest second parental lineage additional to the consensus backbone. Uncertainty regarding the status of a putative *Duparquetia* WGD is marked as a broken arrow reflecting the allopolyploid hypothesis identified in our results. Branch length does not reflect any putative chronological assessments.

## Discussion

There is strong support for multiple types of independent WGDs affecting the legume backbone. Papilionoideae and Detarioideae are autopolyploid. Dialioideae and Cercidoideae are resolved here as clades that are predominantly allopolyploid, but also contain living descendants of a parental lineage diploid relative to the legume ancestor. We also resolve Caesalpinioideae as allopolyploid, fitting several independent lines of reasoning; the specific allopolyploid model depicted is that returned in the reticulate analysis, where a Papilionoideae ancestor hybridized with an extinct lineage sister to all other legumes. *Duparquetia* is depicted as, potentially, an allopolyploid hybrid between two early lineages along the legume backbone — but apart from our assessment that it is polyploid, we have little certainty about the precise origins of any second subgenome in *Duparquetia*, beyond the consensus backbone.

Previous studies have established the non-polyploidy of *Cercis* (5, 6, 12), and the hybrid origin (5, 6) of the non-*Cercis* Cercidoideae, and the results presented here add further support for a single hybridization hypothesis for the origin of all Cercidoideae sister to *Cercis*, with *Cercis* descending from a parental lineage which hybridized with some near-sister lineage to give rise to the remaining Cercidoideae. No convincing evidence was found of a WGD affecting *Cercis*, and direct inference of the likeliest reticulate phylogeny identified allopolyploidy of the non-*Cercis* Cercidoideae in line with previous studies, as the most likely process to account for their duplication status. Our results suggest a similar scenario in Dialioideae; no convincing evidence was found that a WGD affects *Poeppigia*, and our methods identified an allopolyploid model of Dialioideae evolution as the likeliest way of accounting for this disparity in duplication status.

Some of our surprising results should be treated with caution, such as those involving the genera *Poeppigia* and *Duparquetia*. Without genome-scale data for *Poeppigia* equivalent to that available for *Cercis*, the hypothesis of its diploidy relative to the legume ancestor, should be considered preliminary; for now, the data available to us fully accords with it. Relationships between Dialioideae and *Duparquetia* will similarly require better data for clarification. *Duparquetia* had notably higher site concordance with Dialioideae than it did with either Papilionoideae or Caesalpinioideae; yet its consensus phylogenetic position subtending ((PAP, CAE), DIA) would suggest that it is in fact no more closely related to Dialioideae than to the other two.

Multiple complementary lines of evidence in our results suggest that Caesalpinioideae descends from a hybrid between a Papilionoideae ancestor and some other lineage. Gene count data indicates that Caesalpinioideae has clearly undergone a WGD, yet with unusual frequency (84%), paralogous duplication clades of Caesalpinioideae genes seem to be more closely related to distinct extrasubfamilial ortholog clades, especially Papilionoideae ortholog clades, rather than to their own intra-subfamilial paralogs. No evidence was found for models of an additional WGD shared by both Caesalpinioideae and another lineage such as Papilionoideae or Dialioideae; instead, an extinct lineage was inferred as the likeliest alternate parent under the same modeling conditions as above (Fig. 4a, b).

This result casts a new light upon longstanding difficulties in resolving the Caesalpinioideae WGD. While we see no evidence for the multiple rounds of whole genome duplication suggested by Zhao et al. (4), the Caesalpinioideae allopolyploid model does provide an alternative explanation of their observations. Every time an allopolyploid lineage is created, the resulting homeologous chromosomes are related to one another not at the moment of hybridization, but at the speciation node of their hybrid parents. Methodologies based on counting “bursts” of gene duplications, upon encountering a hybrid lineage therefore must see a “burst” of gene duplications taking place at *that* speciation node, the one predating the hybrid’s own origin, belonging to the hybrid’s parents. When PhyloNet infers a lost lineage ancestral to all legumes, as the second parent of the Caesalpinioideae (and perhaps also of *Duparquetia*) (i.e., Fig. 4a, b), this is an inference that the two subgenomes of Caesalpinioideae separated from one another prior to the legume crown node, at the unidentified node where the lost lineage speciated from the rest. Our model thus necessarily contains Zhao et al.’s (4) observed burst of gene duplications at the legume crown node, by observing the same reticulation at the base of Caesalpinioideae, as reported by Koenen et al. (3)

But the consequences of hybridization do not end at the moment of hybridization. Hybridization triggers numerous genome-scale changes (13). Among these are exchanges between homeologous chromosomes, which can lead to widespread paralog substitution. For example, a study of synthetic peanut allopolyploids observed substitution of entire paralogous chromosomes (16). Any sufficiently-extensive episode of paralog substitution occurring in a novel hybrid species will generate a second apparent burst of gene duplications after the node at which the hybrid diverges from its parental lineage, as a result of this paralog replacement. Our model thus necessarily contains both Koenen et al.’s (3) specific observation that Caesalpinioideae failed to collapse to a single lineage, and Zhao et al’s (4) specific observation of a second burst of gene duplication at the crown node of Caesalpinioideae. Hybrids truly bear genomes from multiple lineages, and the interrelationships among the individual genes are easily complicated.

Another seemingly-paradoxical result is the closer-than-expected association between Dialioideae and *Duparquetia* in terms of site concordance fators. The phylogenetic position of *Duparquetia* would ordinarily indicate that the genus ought not be any likelier to bear site similarities with Dialioideae, than with Papilionoideae or Caesalpinioideae. When Dialioideae resolves in gCFs as more closely related to Papilionoideae-Caesalpinioideae, than to *Duparquetia*, despite sCFs more commonly backing a terminal DIA-DUP relationship than DIA-CAE, DIA-PAP, etc., this implies that the gene-tree construction models are positing a PAPCAE ancestor that bears more similarities to the Dialioideae ancestor, than modern *Duparquetia* does. While a nuanced interpretation, there is no direct conflict in this. What it does suggest, is that if further studies firmly establish *Poeppigia* as unduplicated, then as a member of Dialioideae, it may be useful for resolving the most troublesome branches of the legume backbone, in a way similar to *Cercis*.

Given that polyploidy is a major force in plant evolution and speciation (17), the complex reticulation observed in backbone subfamily relationships in Leguminosae, is no great surprise. Complex polyploid networks are still known in the legumes today; a particularly well-studied one is the perennial soybean polyploid complex (18). The K-Pg boundary period when legumes were evolving is well-known as a period when many hybrid lineages survived (19), so as noted by Soltis et al. (20): “The question is no longer ‘What proportion of angiosperms are polyploid?’, but ‘How many episodes of polyploidy characterize any given lineage?’” Polyploidy immediately preceding clade diversification has marked the evolutionary history of numerous clades of comparable importance to the Fabaceae, including Poaceae (21), Brassicaceae (22), and Solanaceae (23); indeed, these WGDs are credited with contributing to the dramatic increase in species richness of these lineages (20).

In a hybridization event, genes that have already been evolving separately for potentially millions of years become sister chromosomes. Yet they remain related to one another not at the moment of hybridization, but at the speciation node of the hybrid’s parents. Methodologies based on counts of bursts of duplicated genes, can therefore be fundamentally vulnerable to incorrect and contradictory conclusions about the number and timing of WGD events, whenever the species under study are allopolyploids (or potential paleoallopolyploids).

Conversely, allopolyploidy neatly explains why seemingly-contradictory observations could be true at once. Homologs within Caesalpinioideae can be more-closely related to other subfamilies than to one another, even if the lineage does not share a WGD with another lineage, if the homologs in question originated as homeologs. Since the multi-step and interrelated processes of hybridization and diploidization can retroactively convert a speciation event that occurred a million years ago, into an origin point for a hybrid’s paralogs, homeologs within Caesalpinioideae can have originated prior to the separation of Caesalpinioideae from unduplicated lineages such as *Cercis*. Our assessment of the legume backbone in that light, resolves previous contradictory findings by concluding that at least three legume subfamilies are allopolyploid relative to the ur-legume, such that reticulation along the backbone is a fundamental fact of legume subfamilial evolution.

## ACKNOWLEDGMENTS

This work was supported by the United States Department of Agriculture, Agricultural Research Service (USDA-ARS) CRIS Project 5030-21000-071-000D. The USDA is an equal opportunity provider and employer. Mention of trade names or commercial products in this article is solely for the purpose of providing specific information and does not imply recommendation or endorsement by the U.S. Department of Agriculture.

This research used resources provided by the SCINet project and/or the AI Center of Excellence of the USDA Agricultural Research Service, ARS project numbers 0201-88888-003-000D and 0201-88888-002-000D.

This work was supported by Natural Sciences and Engineering Research Council of Canada (grants to AB and WC-M

## Notes

### Competing Interest Statement

The authors have declared no competing interest.

